# A Drosophila Circuit for Habituation Override

**DOI:** 10.1101/2021.09.11.459900

**Authors:** Swati Trisal, Marcia Aranha, Ankita Chodankar, K. VijayRaghavan, Mani Ramaswami

## Abstract

Habituated animals retain a latent capacity for robust engagement with familiar stimuli. In most instances, the ability to override habituation is best explained by postulating that habituation arises from the potentiation of inhibitory inputs onto stimulus-encoding assemblies and that habituation override occurs through disinhibition. Previous work has shown that inhibitory plasticity contributes to specific forms of olfactory and gustatory habituation in *Drosophila*. Here we analyze how exposure to a novel stimulus causes override of gustatory (proboscis-extension reflex or “PER”) habituation. While brief sucrose contact with tarsal hairs causes naïve *Drosophila* to extend their proboscis, persistent exposure reduces PER to subsequent sucrose stimuli. We show that in so habituated animals, either brief exposure of the proboscis to yeast or direct thermogenetic activation of sensory neurons restores PER response to tarsal sucrose stimulation. Similar override of PER habituation can also be induced by brief thermogenetic activation of a population of TH (Tyrosine-Hydroxylase) positive neurons, a subset of which send projections to the subesophagial zone (SEZ). Significantly, sensory-neuron induced habituation override requires transmitter release from these TH-positive cells. Treatments that cause override specifically influence the habituated state, with no effect on the naïve sucrose response across a range of concentrations. Taken together with other findings, these observations in female flies are consistent with a model in which novel taste stimuli trigger activity in dopaminergic neurons which, directly or indirectly, inhibit GABAergic cells that drive PER habituation. The implications of these findings for general mechanisms of attentional and sensory override of habituation are discussed.

## INTRODUCTION

Habituation is a form of non-associative learning in which the response to a stimulus reduces after repeated or extended passive exposure. However, a latent ability to respond to the innocuous stimulus remains. “Dishabituation” is a classical, defining feature of the habituated state, distinguishing it from sensory or synaptic fatigue (Thompson and Spencer, 1966; Rankin et al., 2009; Ramaswami, 2014). In this article we often use the term “override” in place of dishabituation, which while largely synonymous, better acknowledges that behavioral responses can be reinstated by not only classic dishabituating (novel) stimuli, but also top-down attentional mechanisms (Cooke and Ramaswami, 2020). For instance, in volitional inspection of a familiar object on a shelf, habituation is overcome by attention to the object, and not by an extraneous novel stimulus.

Habituation and the associated phenomenon of dishabituation/ override are well known across different phyla of animal kingdom (Glanzman et al., 1989; Zaccardi et al., 2004; Smith et al., 2009; Ramaswami, 2014). Two broad observations are relevant here. First, mechanisms underlying very short and longer lasting forms of habituation may differ; some forms of the latter are known to involve inhibitory potentiation (Ramaswami, 2014; Shen et al., 2020). Second, largely due to the difficulty of necessary behavioral experiments, in most (but not all e.g. Semelidou et al., 2018) instances, habituation override is not distinguished from a potential confounding process of response sensitization (Castellucci et al., 1970; Hawkins et al., 1998; Asztalos et al., 2007a; Asztalos et al., 2007b). Thus, although arguments and evidence support a disinhibitory mechanism (Fischer et al., 1997; Das et al., 2011; Kato et al., 2015; Ogg et al., 2018) the neural pathways and mechanisms of habituation override remain incompletely characterized. Here we address this issue in the gustatory system of *Drosophila*.

*Drosophila* sample and taste potential foods via chemosensory hairs on their tarsi and their proboscis (Stocker, 1994; Montell, 2009). Sugars detected by sensory hairs trigger a proboscis extension reflex (PER), which enables feeding (Minnich, 1921; Dethier, 1976). The neural circuit for PER is only partially understood. Sensory information is processed in subesophagial zone (SEZ) and then communicated to command neurons whose activation triggers the motor programme required for PER (Flood et al., 2013). Repeated sucrose stimulation of the tarsus under conditions where proboscis extension is futile, leads to reduced PER response through a process that shows several classic features of habituation (Duerr and Quinn, 1982; Le Bourg, 1983; Fois et al., 1991; Engel and Wu, 2009; Paranjpe et al., 2012). Importantly, in habituated animals, PER to sucrose is quickly restored if the fly is presented with a strong, novel sensory stimulus (Le Bourg, 1983; Fois et al., 1991; Paranjpe et al., 2012). Here, we investigate circuit mechanisms that drive override of PER habituation.

We first reproduced prior experiments providing key support for increased inhibition in the PER pathway being the core mechanism for PER habituation (Paranjpe et al., 2012). Thereafter, we addressed mechanisms of habituation override, which we found could be achieved by yeast stimulation of the proboscis, thermogenetic activation of yeast-responsive or bitter responsive sensory neurons, or activation of a dopaminergic neuron subpopulation. We show that sensory stimulation procedures induce habituation override through dopaminergic neuron activation. Crucially, each dishabituation protocol specifically affects the response of habituated animals; none sensitize the naïve response. While the data do not yet conclusively define all elements of the dishabituation circuit or the mechanism by which dopaminergic neurons trigger override, they support a model in which novel stimuli induce dopamine release, which acts to directly or indirectly inhibit inhibitory neurons that drive PER habituation. We suggest that this work: (a) circumscribes core elements of a sensory-central circuit for habituation override; (b) provides evidence for a new disinhibitory pathway in the Drosophila brain; and (c) supports an emerging framework in which latent perceptions, memories and behaviors may be generally activated through disinhibition (Sridharan and Knudsen, 2015; Barron et al., 2017; Wang and Yang, 2018).

## MATERIALS AND METHODS

### Experimental Design

#### Drosophila stocks

Fly stocks were maintained on standard corn meal media. Canton S (CS) flies were used as wild-type controls unless otherwise stated. The stocks were obtained either from stock centres or as generous gifts from following sources: *Ir25a-Gal4* was provided by Carlos Ribeiro (Champalimaud Centre for the Unknown, Lisbon, Portugal), *TH-C’-Gal4, TH-D’-Gal4* and *TH-C-Gal80* were generously provided by Mark Wu (Johns Hopkins University, Baltimore, MD), *Gad1-Gal4* was from Gero Miesenbock (Oxford University, Oxford, UK), *TH-Gal4* was provided by Gaiti Hassan (National Centre for Biological Sciences, Bangalore, India), *Gr66a-LexA, lexAop-CD4::spGFP11; UAS-CD4::spGFP1-10* and *LexAop-TRPA1* were provided by Kristin Scott (University of California, Berkley). *rut^2080^* and *UAS-rut^+^* were provided by Martin Heisenberg. *UAS-Shi^ts^* was obtained from Toshi Kitamoto (University of Iowa, Iowa city, IA), *UAS-TRPA1* was provided by Paul Garrity (Brandeis University, Waltham, MA). *Wg/CyO; Gr66a-Gal4* (BL 57670), *UAS-mCD8::GFP* (BL 5130), *UAS-CD8::RFP*, *LexAop-CD8::GFP* (BL 32229) were obtained from Bloomington Drosophila Stock Centre. Dopamine receptors were downregulated using lines *UAS-DopR1-miR, UAS-DopR2-miR* and *UAS-D2R-miR* obtained from Mark Wu (Johns Hopkins University, Baltimore, MD) in *Gad1-Gal4* expressing neurons (data not shown). Tyrosine hydroxylase was downregulated in *TH Gal4* neurons using TH-miR-G/TM6, Sb and TH-miR-9/Cyo; TH-miR-2/TM6, Tb provided by Mark Wu (Johns Hopkins University, Baltimore, MD) (data not shown). However, the results were inconclusive.

#### Proboscis extension behaviour

Proboscis extension behaviour was carried out as described in (Paranjpe et al., 2012). Briefly, 3-4 days old female flies were mounted individually, ventral side up, on a cover slip. The flies were kept in a humidified chamber for 2 hours for recovery. Tarsal hairs were stimulated with 2% sucrose using 1ml syringe needle. The naïve response score was determined after stimulating tarsal hairs five times. Failed PER response was scored as zero and complete proboscis extension was scored as one. Flies that showed naïve PER response less than three out of five times were discarded and not used for habituation experiments.

### Habituation of proboscis extension reflex

Tarsal hairs were stimulated with 10% sucrose for 10 minutes after which the legs were washed using distilled water. The habituated response or post exposure response was recorded by stimulating tarsus with 2% sucrose five times, similar to naïve response.

### Habituation Override

10% yeast was presented to the proboscis over one minute, in a “spaced manner” to prevent consumption of yeast and satiation. In gist, the yeast paste was applied to labellar bristles for about a second but withdrawn rapidly before consumption could occur and this was repeated (about 6 times) over a one minute period. After exposure to yeast, flies were allowed to groom for 1 minute so that yeast sticking to the proboscis could be cleaned. PER responses were subsequently recorded as usual.

For habituation override using mechanical stimulus, flies mounted on the cover slip were placed in a petri dish (35mm) and vortexed for 1 minute on a Scientific Industries Vortex-Genie 2 instrument at the highest setting. Flies were allowed to recover from the shock for 1 minute before testing the response.

To distinguish override from sensitization, naïve response of flies was recorded at concentrations of sucrose lower than 2%. This was done as the PER response is maximum at 2%, any difference as a result of manipulation would have been difficult to observe. Four concentrations of sucrose were tested- 0.1%, 0.5%, 1% and 2%. Flies were then exposed to either stimulus used for override or heat, for 1 minute. Post response was tested after 1 minute rest at RT.

### Heat-mediated manipulation

For TRPA1 and Shi^ts^ experiments, flies were transferred to 32° and 34°C respectively on a dry bath for 1 minute. In case of Shi^ts^ experiments, proboscis was stimulated with 10% yeast while at 34°C. Flies were given 1 minute rest at RT before testing PER response.

### Immunohistochemistry

Adult brains were dissected in 1X PBS and fixed in 4% paraformaldehyde diluted in 1X PBS with 0.3% Triton-X (PTX) for 30 minutes at room temperature. Samples were washed with 0.3% PTX for 20 minutes three times at room temperature and incubated with primary antibody for 48 hours at 4°C on a shaker. Samples were again washed three times with 0.3% PTX for 20 minutes each. Secondary antibodies conjugated with Alexa Fluor-488, Alexa Fluor-568 and Alexa Fluor-647 (1:500, Invitrogen), diluted in 0.3% PTX, were added and samples were incubated for 24 hours. Brain samples were again washed repeatedly three times for 20 minutes each and mounted in Vectashield (H-1000, Vector laboratories) on a glass slide with spacers. Images were acquired using Zeiss LSM 510 Meta microscope and Olympus Fluoview (FV-3000).

Following primary antibodies were used: rabbit anti-GFP (1:100, invitrogen), mouse anti-Bruchpilot (n82) (1:20, DSHB), mouse anti-GFP (Sigma, 1:100), rabbit anti-TH (1:1000, Invitrogen), chick anti-GFP (Abcam, 1:5000), rabbit anti-DsRed (1:100, Clontech).

### Statistical analysis

Non parametric tests were used to analyse data. Friedman test was used to analyse habituation override experiments which had three matched groups, followed by Dunn’s multiple comparisons post hoc test to determine difference among different treatments. Wilcoxon sign rank test was used for comparing two paired groups whereas Mann Whitney U test was used to compare two unpaired groups in Fig1C. All the data was analysed using GraphPad Prism v8.4.2 software.

**Figure 1.**
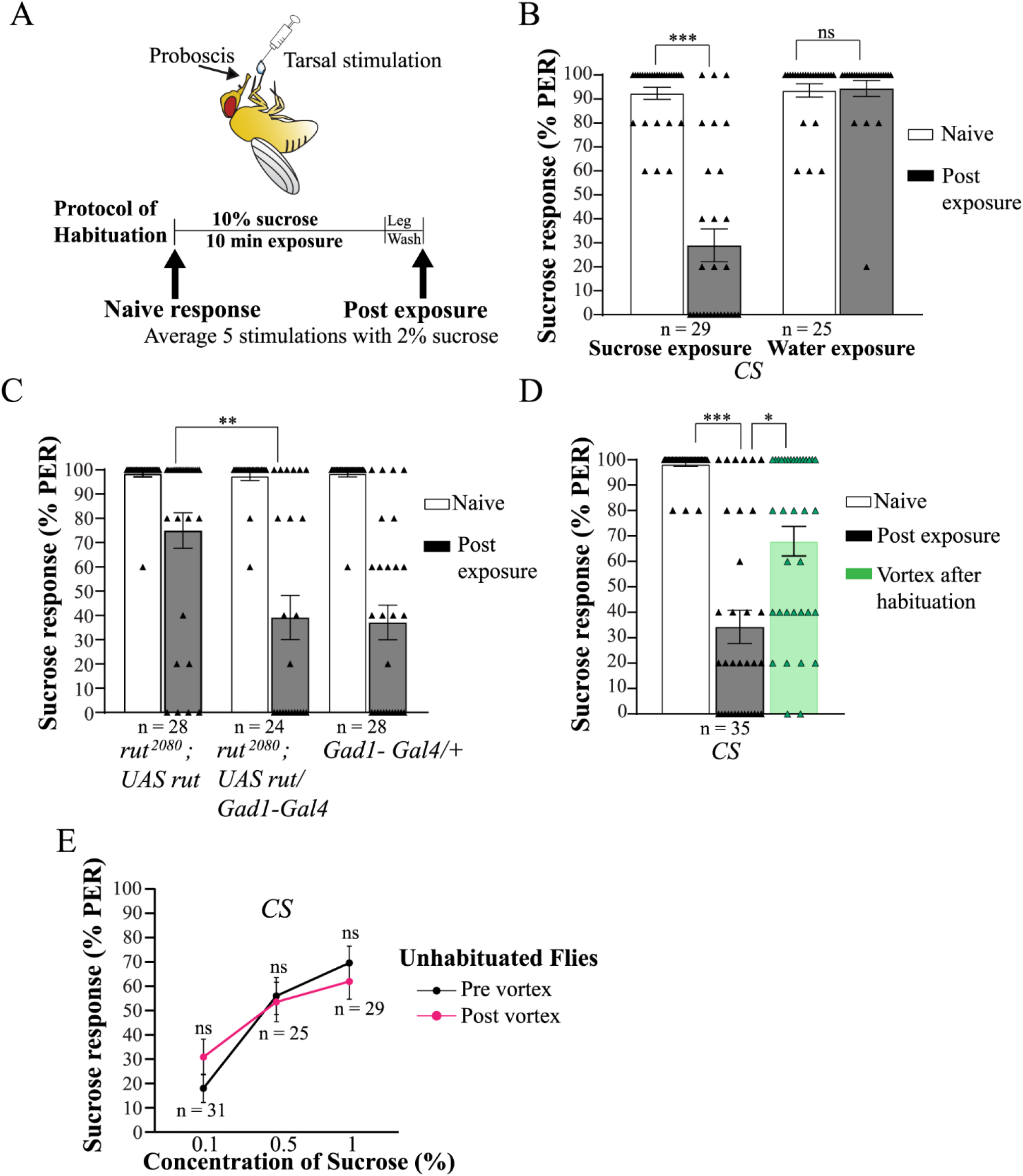
Gustatory habituation of PER to sweet taste. **(A)** Behavioural protocol of PER habituation to tarsal stimulation following sucrose exposure. **(B)** Tarsal exposure to 10% sucrose for 10 minutes leads to decrease in PER response (***p< 0.0001, Wilcoxon signed rank test) whereas exposure to water does not affect the response (p = 0.6133, Wilcoxon signed rank test). **(C)** As previously described (Paranjpe et al, 2012), gustatory habituation is dependent on the Rutabaga adenylate cyclase since it is impaired in *rut^2080^* mutants. Habituation can be restored by expressing wild type *rut* in inhibitory *Gad1-Gal4* neurons (**p < 0.006, Mann Whitney U stat = 227). **(D)** The habituated response can be overridden by a novel mechanical stimulus like vortexing (p<0.001, F-stat = 47.08, Dunn’s multiple comparison post-hoc test, ***p <0.001, *p = 0.01). **(E)** Vortexing does not have an effect on naïve response to sucrose at concentrations 0.1% (p = 0.0518, Wilcoxon signed rank test), 0.5% (p= 0.6880, Wilcoxon signed rank test) and 1 % (p= 0.1626, Wilcoxon signed rank test) as, determined by testing PER response pre and post vortexing at different sucrose concentrations. Therefore, the enhanced PER response after vortexing habituated animals is not a result of general sensitization but instead represents a specific override of habituation to sucrose. Bars represent mean±SEM, triangles represent individual data points, ns represents not statistically significant, p>0.05.

## RESULTS

### Habituation of the proboscis-extension reflex to sucrose

Two-percent sucrose applied to tarsal hairs of immobilized, naïve flies induces robust and reproducible proboscis extension response. However, following extended 10-minute exposure to 10% sucrose solution, PER decreases from 92.41% to 28.96% (***p<0.0001, Wilcoxon test) (Fig 1A-B). No change in naïve PER response is seen if water is presented for 10 minutes instead of 10% sucrose (Fig1B). A previous study concluded that plasticity in central GABAergic neurons is required for PER habituation (Paranjpe et al., 2012). Given their significance for the inferred inhibitory mechanism for PER habituation, and because these observations were made exclusively in male flies, we independently repeated key experiments to re-examine: first, the need for the *rutabaga*-encoded adenyl cyclase in PER habituation; and second, its reported sufficiency in *GAD1-Gal4* expressing, predominantly GABAergic neurons for this function in our hands and in female flies. Our results were consistent with and confirmed previously reported observations (Figure 1C).

Habituation is often distinguished from sensory or muscular fatigue by demonstrating the rapid reinstatement of the sensory response by strong, novel stimuli (Rankin et al., 2009; Ramaswami, 2014). Consistently, PER-habituated animals retain the ability to respond relatively robustly to sucrose, as evidenced following strong or novel sensory stimulation. Similar to olfactory habituation, strong mechanical stimulation (vortexing in a dish) for 1 minute substantially reinstates the sucrose-induced PER in habituated flies (p< 0.001, F-stat = 47.08, Dunn’s multiple comparisons post-hoc test, *p= 0.01), without affecting baseline sucrose sensitivity in naïve animals (Figure 1D-E; (Das et al., 2011; Paranjpe et al., 2012). Thus, the effect of mechanical stimulation on PER is selective on the habituated state and represents a form of habitation override and not sensitization. In order to address the underlying circuit mechanisms, we first looked to identify more precisely defined sensory stimuli that could cause override of PER habituation.

### Novel stimuli override PER habituation

*Drosophila* are attracted by the taste of yeast. Gustatory receptor neurons (GRNs) that respond to yeast components have been recently identified (Fischler et al., 2007; Wisotsky et al., 2011; Ganguly et al., 2017); (Steck et al., 2018). To examine whether the novel taste of yeast could influence PER habituation to sucrose, we applied a 10% yeast solution to the fly labellum, which has previously been shown to possess yeast-responsive GRNs (Steck et al., 2018), and tested if this resulted in override of sucrose habituation. Application of yeast to the labellum substantially reinstated the PER response to tarsal sucrose stimulation in PER-habituated animals (p<0.0001, F-stat = 31.15, Dunn’s multiple comparison post-hoc test, *p= 0.014; Fig 2A-B). In contrast, 10% sucrose solution, which should be familiar to the flies, when applied to the labellum of habituated flies had no effect on PER (Fig 2B). Thus, a brief experience of a novel and in this case, attractive stimulus appears capable of inducing habituation override. We further looked to identify a single GRN class responsive to yeast components that may be sufficient to drive dishabituation. To do this, we expressed heat-activated cation-permeable TRPA1 channels in yeast-responsive GRNs and tested if heat-induced activity in these cells, which bypassed the need for normal ligand-receptor interactions, would result in dishabituation (Hamada et al., 2008). Such experiments showed that “thermogenetic” activation of the Ir25a class of sensory neurons is sufficient to cause PER habituation override.

**Figure 2.**
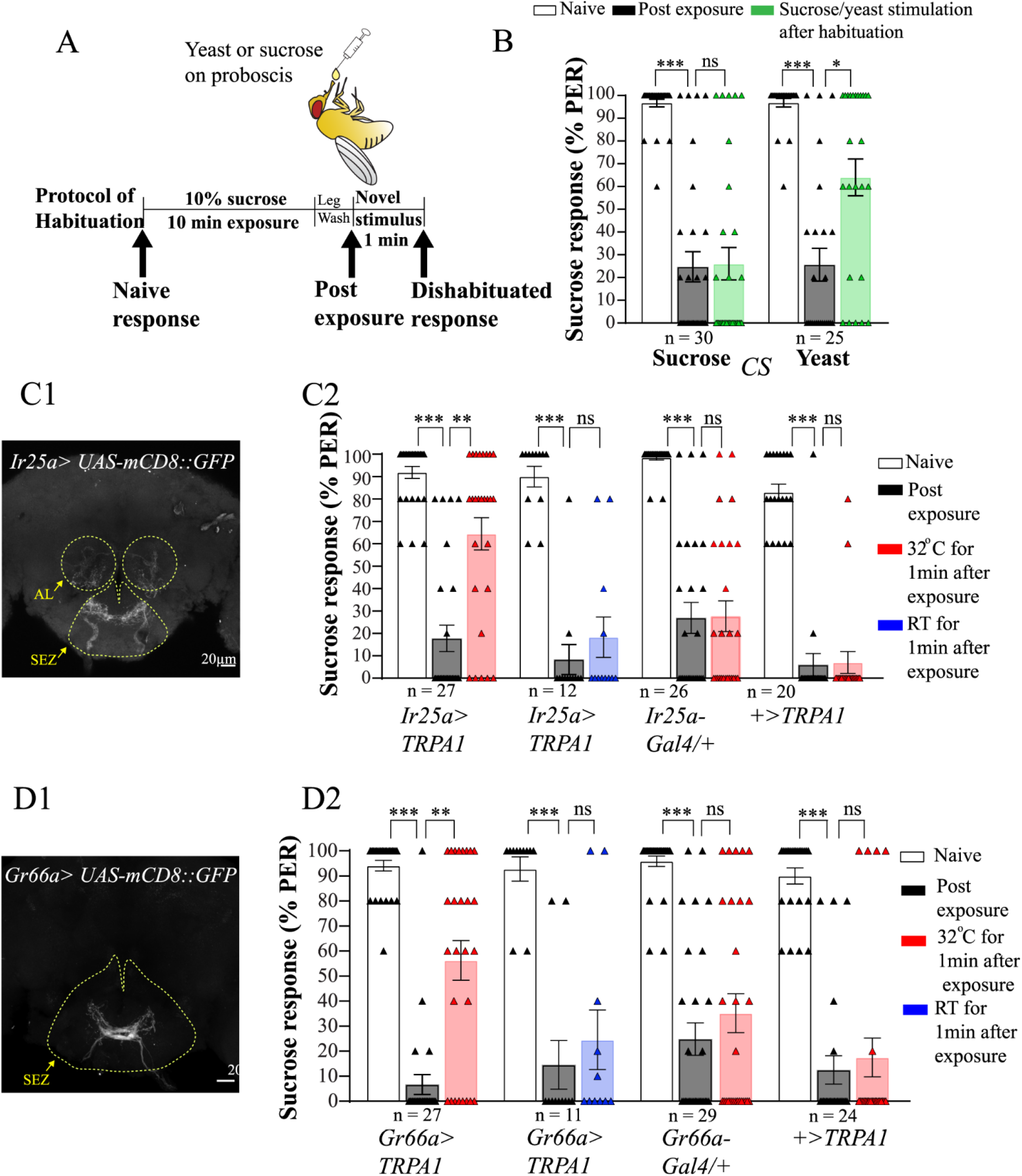
Habituation override mediated by novel stimulus. **(A**) Behaviour protocol of PER habituation and its override by presenting stimulus to the labellum **(B)** Presentation of novel 10% yeast stimulus to labellum restores PER response (p<0.0001, F-stat = 31.15, Dunn’s multiple comparisons post-hoc test, ***p<0.0001, *p = 0.014) whereas 10% sucrose, which is a familiar stimulus, does not have an effect on habituated response (p<0.0001, F-stat = 43.49, Dunn’s multiple comparisons post-hoc test, ***p<0.0001, p > 0.99). **(C1)** Expression pattern of *Ir25a-Gal4* in the adult brain. Ir25a-Gal4 is expresssed in gustatory receptor neurons, predominantly bitter and sweet sensory neurons in the subesophageal zone (SEZ), as well as some olfactory receptor neurons in the antennal lobe (AL). **(C2)** Directly activating yeast responsive *Ir25a-Gal4* cells expressing heat activated channel *TRPA1* for 1 minute after habituation at 32°C, is sufficient to override PER habituation to sucrose (p<0.0001, F-stat = 38.66, Dunn’s multiple comparisons post-hoc test, ***p<0.0001, **p = 0.0012) whereas at permissive temperature, *Ir25a>TRPA1* flies do not show an increase in habituated response (p<0.0001, F-stat = 22.62, Dunn’s multiple comparisons post hoc test, ***p = 0.0002, p>0.99) (*Ir25a-Gal4/+* p<0.0001, F-statistic = 38.99, Dunn’s multiple comparison post-hoc test, ***p<0.0001, p > 0.99, *UAS-TRPA1/+*, p<0.0001, F-stat = 33.88, Dunn’s multiple comparisons post-hoc test, ***p<0.0001, p > 0.99. **(D1)** Expression pattern of *Gr66a-Gal4* in the subesophageal zone (SEZ) region of adult brain. The bitter sensory neurons marked by Gr66a form a ringed structure; they project to the anterior region of the SEZ. **(D2)** Thermogenetic activation of bitter *Gr66a-*marked GRNs for 1 minute after habituation at 32°C, also overrides sweet-taste habituation (p<0.0001, F-stat = 39.85, Dunn’s multiple comparisons post-hoc test, ***p<0.0001, **p = 0.0042) whereas no difference is observed in controls (Gr66a>UAS-TRPA1 at RT p<0.0001, F-stat = 17.56, Dunn’s multiple comparisons post hoc test, ***p= 0.0009, p>0.99, *Gr66a-Gal4/+* p<0.0001, F-stat = 44.60, Dunn’s multiple comparisons post-hoc test, ***p<0.0001, p = 0.97, *UAS-TRPA1/+*, p<0.0001, F-stat = 40.22, Dunn’s multiple comparison post-hoc test, p<0.0001, p > 0.99). Together the data show that stimulus-novelty drives override of habituation. Bars represent mean±SEM, triangles represent individual data points, ns represents not statistically significant, p>0.05.

**Figure 3.**
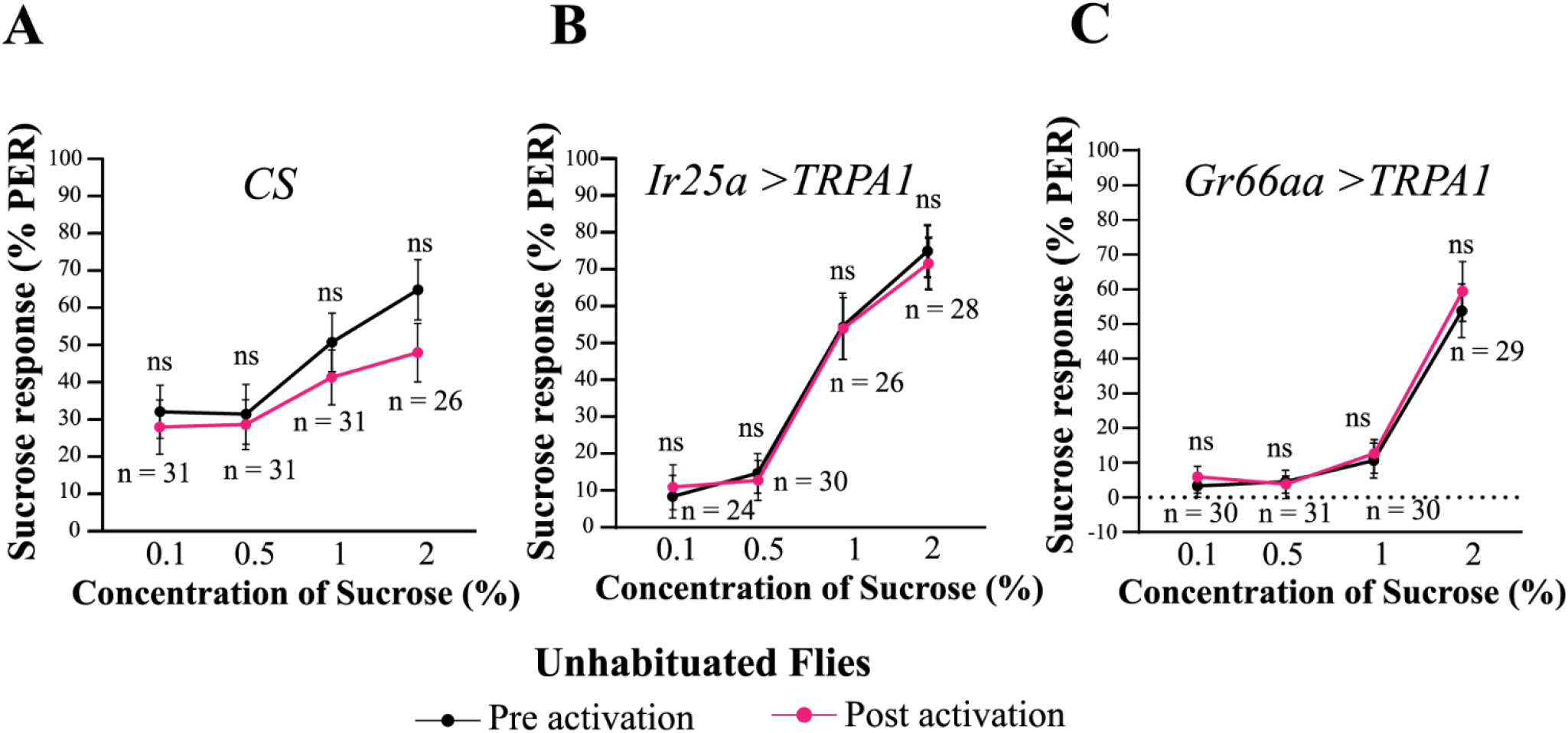
Novelty induced habituation override does not induce sensitization of sensory response. **(A)** Labellar exposure to 10% yeast does not increase the naïve PER response to 0.1%, 0.5%, 1% or 2% sucrose stimulation (Probabilities of PER being unchanged using the Wilcoxon sign rank test: p = 0.6404, p = 0.6731, p = 0.1957, and p = 0.1208 at the 4 respective sucrose concentrations). **(B)** Thermogenetic activation of *Ir25a-Gal4* does not sensitize response to sucrose tested at concentration 0.1% (p > 0.99, Wilcoxon sign rank test), 0.5% (p = 0.5938, Wilcoxon sign rank test), 1% (p = 0.7588, Wilcoxon sign rank test), 2% (p = 0.4814, Wilcoxon sign rank test,). **(C)**Thermogenetic activation of *Gr66a-Gal4* does not have an effect on response to 0.1% (p = 0.5, Wilcoxon sign rank test), 0.5% (p > 0.99, Wilcoxon sign rank test), 1% (p = 0.375, Wilcoxon sign rank test), 2% (p = 0.3828, Wilcoxon sign rank test) concentrations of sucrose. Points represent mean±SEM, ns represents not statistically significant, p>0.05.

TRPA1 channels open at temperatures above 25°C; thus, activation of TRPA1-expressing neurons can be temporally controlled by exposing experimental animals to temperatures where the channel is either closed (RT) or open (above 25°C). We expressed TRPA1 in yeast responsive *Ir25a-Gal4* positive sensory neurons (Steck et al., 2018). These flies were habituated to sucrose at room temperature (RT, 21°C), transferred to 32°C post-habituation for 1 minute and then tested for PER. Thermogenetic activation of Ir25a-expressing GRNs was sufficient to cause rapid override of sucrose habituation (Fig 2C). Thus, after Ir25a activation, PER-habituated animals showed significantly increased PER to tarsal sucrose stimulation (p <0.0001, F-stat = 38.66, Dunn’s multiple comparison post-hoc test, **p=0.0012). Similarly, 32°C exposure of genetic control animals not expressing TRPA1 did not affect PER habituation (Fig 2C).

To distinguish between the novelty and attractiveness of yeast taste as being primarily instrumental in override, we also examined, whether thermogenetic activation of bitter-compound responsive Gr66a expressing GRNs could similarly affect override of PER habituation. Thus, we expressed TRPA1 in *Gr66a-Gal4* positive neurons and examined how sucrose-responsiveness of flies habituated at RT was altered after a brief 1-minute shift to 32°C. Sucrose induced PER increased from 6.67% to 56.29% (p<0.0001, F-stat = 39.85, Dunn’s multiple comparison post-hoc test, **p = 0.0042) indicating that brief activation of bitter-taste sensing *Gr66a*-positive neurons is sufficient to induce dishabituation. The observation that perception of novel bitter taste as well as novel yeast taste can promote override of PER habituation strongly indicates that it is the novelty of the dishabituating stimulus, rather than its attractiveness, that drives habituation override (Fig 2D). We also tested the effect of thermogenetic activation of olfactory and visual sensory neurons by expressing TRPA1 in all *Or83b-Gal4* positive olfactory sensory neurons (OSNs), in subsets (*Or85a-Gal4* or *Gr21-Gal4* positive) of OSNs that mediate strong repulsion, or *Rh1-Gal4* expressing photoreceptors, but we did not observe an effect on PER habituation (data not shown). Therefore, we did not explore these stimuli further.

Does novelty-induced override of PER habituation represent a specific effect on the habituated state or a general sensitization of the sensory response? To differentiate between these possibilities, we tested whether and how yeast exposure, *Ir25a-*neuron activation or *Gr66a*-neuron activation altered PER in naïve flies to a range of sucrose concentrations. If the increase in PER response in habituated flies were due to sensitization, then we would expect an increase in PER after the novel stimulus is applied to naïve animals. However, as shown in 3A, 3B and 3C, there is no significant difference in PER response before and after presentation of yeast or *Ir25a* or *Gr66a*-GRN activation respectively. Thus, it can be concluded that the reinstatement of PER response that we observe in our experiments is due to a process that specifically acts on neural correlates of PER habituation, not broadly on taste perception.

### Artificial activation of Tyrosine-Hydroxylase (TH) expressing neurons overrides PER habituation

Across species, novel stimuli trigger central release of neuromodulators that confer or enhance their salience (Ranganath and Rainer, 2003; Kafkas and Montaldi, 2018). In particular, the activity of dopaminergic neurons has been implicated in the novelty response both in insects and in mammalian systems, (Hattori et al., 2017; Morrens et al., 2020). We therefore investigated whether thermogenetic activation of *Drosophila* tyrosine-hydroxylase (TH) expressing central dopaminergic neurons would be (a) sufficient and (b) necessary for novel-taste induced override of PER habituation.

In PER-habituated *TH-Gal4>UAS-TRPA1* animals, 1-minute exposure to 32°C to drive thermogenetic activation of *TH-Gal4* expressing neurons resulted in rapid habituation override (Fig 4A). In PER-habituated animals, sucrose-induced PER was close to 12%; in these same animals, activation of TH expressing neurons increased PER to 60% (p<0.0001, F-stat = 50.54, Dunn’s multiple comparisons post-hoc test, ***p = 0.0004). Significantly, activation of TH Gal4 neurons had no effect on the innate response to sucrose in naïve flies (Fig 4B). Thus, activation of these modulatory neurons causes habituation override, not general taste sensitization.

**Figure 4.**
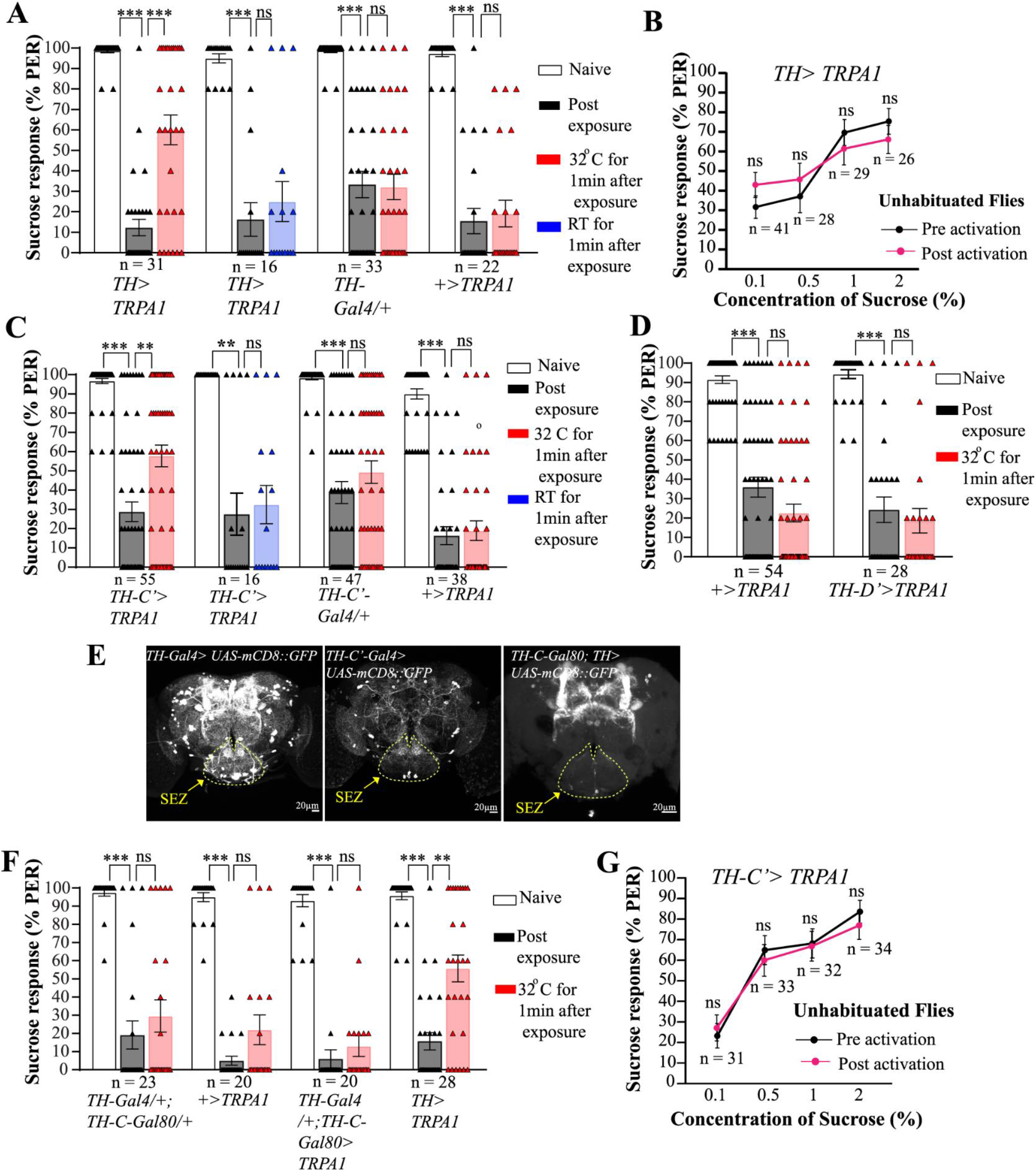
Activation of TH expressing neurons overrides habituation. **(A)** Thermogenetic activation of TH expressing neurons for 1 minute at 32°C after habituation results in habituation override (p<0.0001, F-stat = 50.54, Dunn’s multiple comparisons post-hoc test, ***p<0.0001, ***p = 0.0004) (*TH-Gal4/+*, p<0.0001, F-stat = 49.56, Dunn’s multiple comparisons post-hoc test, ***p<0.0001, p >0.99, *UAS-TRPA1/+*, p <0.0001, F-stat = 37.31, Dunn’s multiple comparisons post-hoc test, ***p<0.0001, p>0.99, *TH-Gal4/UAS-TRPA1* temperature control, p<0.0001, F-stat = 26, Dunn’s multiple comparisons post-hoc test, ***p<0.0001, p = 0.8665). **(B)** Activation of TH expressing neurons does not lead to sensitization of gustatory response tested at 0.1% (p = 0.09, Wilcoxon sign rank test), 0.5% (p = 0.1, Wilcoxon sign rank test), 1% (p = 0.14, Wilcoxon sign rank test), 2% (p = 0.1367, Wilcoxon sign rank test). **(C)** Activation of a subset of TH expressing cells marked by *TH-C’-Gal4* for 1 minute after habitation at 32°C is sufficient to override habituated response to sucrose (p<0.0001, F-stat = 69.50, Dunn’s multiple comparisons post-hoc test, ***p<0.0001, **p = 0.0042) (*TH Gal4/+*, p<0.0001, F-stat = 59.70, Dunn’s multiple comparisons post-hoc test, ***p<0.0001, p = 0.4914 *, UAS-TRPA1/+*, p<0.0001, F-stat = 63.52, Dunn’s multiple comparisons post-hoc test, ***p<0.0001, p > 0.99) (*TH-C’-Gal4* temperature control, p<0.0001, F-stat = 21.41, Dunn’s multiple comparisons post-hoc test, **p = 0.0017, p > 0.99). **(D)** Activation of *TH-D’* subset of neurons, which marks subset of neurons that does not overlap with *TH-C,’* for 1 minute after habituation at 32°C does not override habituation (p<0.0001, F-stat = 42.70, Dunn’s multiple comparisons post-hoc test, ***p <0.0001, p> 0.99) (*UAS-TRPA1/+*, p<0.0001, F-stat = 76.79, Dunn’s multiple comparisons post-hoc test, ***p<0.0001, p = 0.3710). **(E)** Combining *TH-C-Gal80* along with *TH Gal4* blocks the expression in *TH-C’* subset of neurons only, as observed by GFP expression (SEZ represents subesophageal zone). **(F)** The flies carrying *TH-C-Gal80* along with *TH-Gal4* fail to show habituation override when these are activated at 32°C after habituation further confirming the role of *TH-C’* subset of neurons (p<0.0001, F-stat = 35.74, Dunn’s multiple comparisons post-hoc test, ***p<0.0001, p >0.99) (*TH-Gal4/ UAS-TRPA1*, p<0.0001, F-stat = 45.65, Dunn’s multiple comparisons post-hoc test, ***p<0.0001, **p = 0.0025) (F-stat = 35.38, *TH-C-Gal80/+*, p <0.0001, F-stat = 35.38, Dunn’s multiple comparisons post-hoc test, ***p<0.0001, p = 0.9061) (*UAS-TRPA1/+*, p<0.0001, F-stat = 30.55, Dunn’s multiple comparisons post-hoc test, ***p<0.0001, p > 0.99). **(G)** Activation of *TH-C’* Gal4 neurons does not affect the naïve response to sucrose tested at 0.1% (p = 0.5635, Wilcoxon sign rank test), 0.5% (p = 0.3748, Wilcoxon sign rank test), 1% (p = 0.7338, Wilcoxon sign rank test), 2% (p = 0.3125, Wilcoxon sign rank test). Bars represent mean ±SEM, triangles represent individual data points, points in figure 4B and 4G represent mean ±SEM, ns represents not statistically significant, p>0.05.

*TH-Gal4* labels around 200 neurons in the Drosophila brain (Friggi-Grelin et al., 2003). In order to more tightly define *TH-*expressing cells involved in PER-habituation override, we tested two non-overlapping subsets of *TH-Gal4* positive neurons, marked by *TH-D’-Gal4* and *TH-C’-Gal4* drivers (which labels ~54±5 and ~45±3 neurons respectively) for their potential roles. While 1-minute thermogenetic activation of *TH-D’* neurons had no effect on PER habituation, similar activation of neurons labelled by *TH-C’-Gal4* significantly increased PER response from 28.72% in control habituated flies, to 57.81% after *TH-C’* activation (p<0.0001, F-stat = 69.50, Dunn’s multiple comparisons post-hoc test,**p= 0.0042 Fig 4C). Moreover, *TH-Gal4*-driven habituation override required activity in *TH-C’* cell population: thus, 32°C exposure did not trigger PER habituation override in *TH-C-Gal80; TH-Gal4> UAS TRPA1* flies, in which Gal80 expression prevented TRPA1 expression in the *TH-C’* subset of *TH-Gal4* neurons (Fig 4D). This indicates that activity in *TH-C’* subset of cells is necessary and sufficient to cause habituation override. To further support this conclusion, we checked and confirmed that activation of *TH-C’* neurons had no significant effect on the PER of naïve flies across a range of sucrose concentrations (Fig 4E) confirming that *TH-C’* neurons specifically influence mechanisms of habituation, rather than general taste perception.

Significantly, a small subset of *TH-C’* neurons (~13±6) are present locally in the SEZ (suboesophagial zone), an area which not only receives inputs from taste sensory neurons, but also houses interneurons and motor neurons involved in proboscis extension (Gordon and Scott, 2009; Kain and Dahanukar, 2015). Thus, some *TH-C’* neurons are well positioned to mediate sensory-driven novelty signals that influence mechanisms of PER habituation.

### A subset of TH-expressing neurons mediates novelty-induced habituation override

In order to test whether the activity of *TH-Gal4* and *TH-C’-Gal4* neurons is necessary for novel-taste induced override of PER habituation, we examined whether novel-taste stimulation could cause habituation override under conditions where synaptic output from *TH-Gal4* or *TH-C’-Gal4* neurons was blocked. To achieve this, we expressed the temperature-sensitive, dominant-negative Shi^ts1^ mutant form of dynamin in these neurons and tested whether a one-minute exposure to 10% yeast at 34°C could override habituation in *TH-Gal4, UAS-Shi^ts1^* and *TH-C’-Gal4 UAS-Shi^ts1^* at temperatures restrictive for Shi^ts1^ dynamin function.

At permissive (room) temperatures, *TH-Gal4, UAS-Shi^ts1^* and *TH-C’-Gal4, UAS-Shi^ts1^* flies behaved similarly to wild-type flies, showing both robust PER habituation after 10 minutes of tarsal sucrose exposure (Fig 5A) and significant PER dishabituation following brief (1 min) exposure of their labella to 10% yeast solution (Fig 5A). The respective efficiency of PER in habituated and dishabituated animals (before and after yeast exposure) were 34.05% and 77.29% (p<0.0001, F-stat = 42.47, Dunn’s multiple comparisons post-hoc test, ***p = 0.0008) and 13.46% and 48.46% (p<0.0001, F-stat = 41.01, Dunn’s multiple comparisons post-hoc test, *p = 0.045) respectively (Fig 5A).

**Figure 5.**
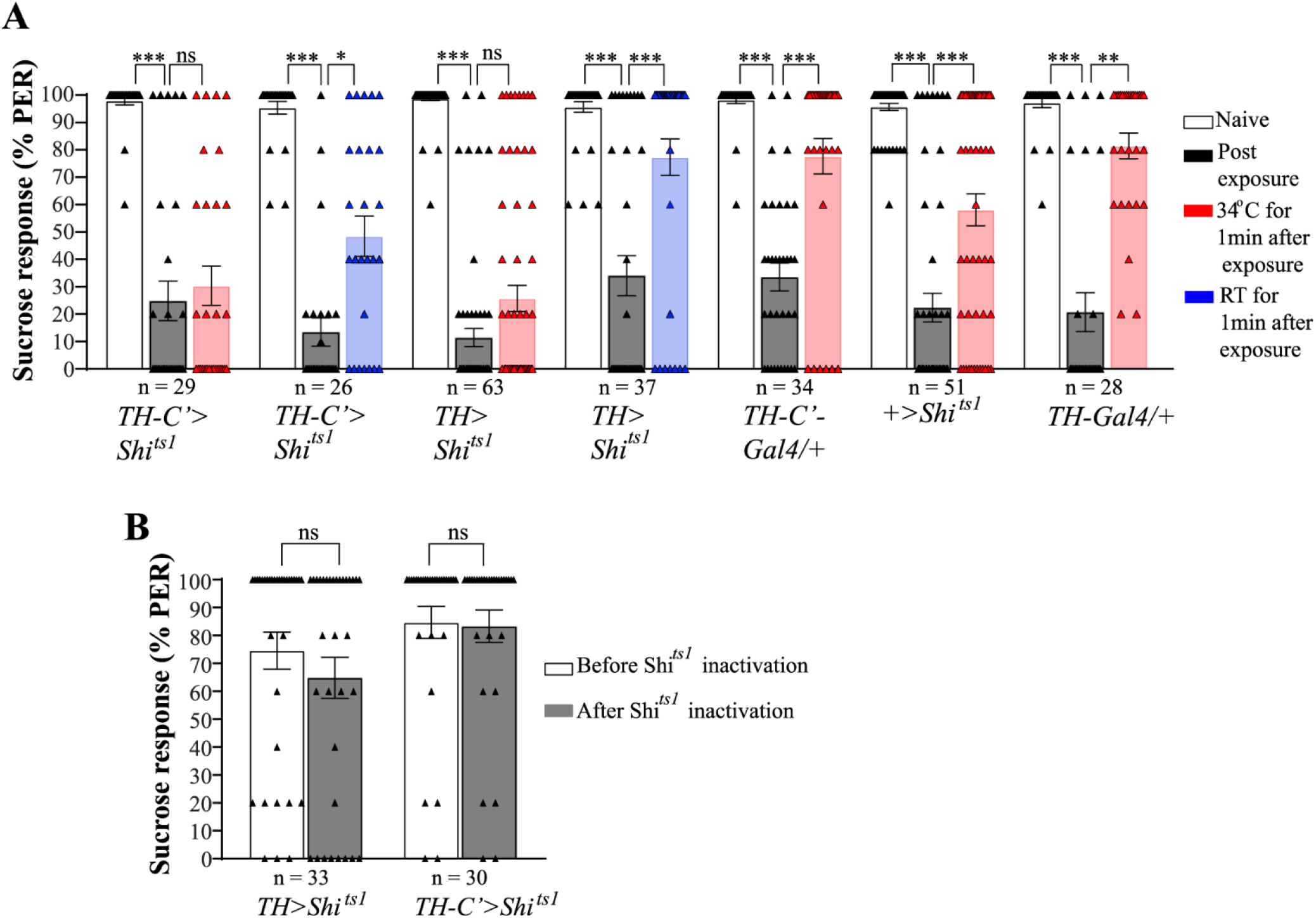
Activity of *TH-C’* subset of neurons is necessary for novelty induced override. **(A**) At permissive temperature (RT = 21°C), presentation of 10% yeast to the labellum overrides habituation in both *TH-C’ Gal4/ UAS-Shit^s1^* (p<0.0001, F-stat = 41.01, Dunn’s multiple comparisons post-hoc test,***p<0.0001, *p = 0.0457) and *TH-Gal4/+;UAS-Shi^ts1^/+* flies and (p<0.0001, F-stat = 42.47, Dunn’s multiple comparisons post-hoc test, ***p<0.0001, ***p = 0.0008) respectively. However, at restrictive temperature (34°C) blocking the synaptic transmission of *TH-Gal4* and *TH-C’-Gal4* neurons by expressing *Shi^ts1^* during presentation of novel yeast stimulus after habituation to sucrose, impairs habituation override (*TH-Gal4/+;UAS-Shi^ts1^/+*, p<0.0001, F-stat = 102.8, Dunn’s multiple comparisons post-hoc test,, ***p<0.0001, p = 0.3897) (*TH-C’-Gal4/UAS-Shi^ts1^*, p<0.0001, F-stat = 37.98, Dunn’s multiple comparisons post-hoc test, ***p<0.0001, p > 0.99) respectively. The genotypic controls show override of habituation to 10% yeast presented to the labellum (*TH-Gal4/+*, p<0.0001, F-stat = 35.83, Dunn’s multiple comparisons post-hoc test, ***p<0.0001, **p = 0.0015) (*UAS-Shi^ts1^/+*, p<0.0001, F-stat = 56.96, Dunn’s multiple comparisons post-hoc test, p<0.0001, ***p = 0.0009), (*TH-C’-Gal4/+*, p<0.0001, F-stat = 46.90, Dunn’s multiple comparisons post-hoc test, ***p<0.0001, ***p = 0.0001) **(B)** Blocking *TH-Gal4* neurons and *TH-C’-Gal4* neurons does not have an effect on PER itself (p = 0.1909 and p = 0.9141, respectively, Wilcoxon sign rank). Bars represent mean ± SEM, triangles represent individual data points, ns represents not statistically significantly, p>0.05.

In contrast, if after PER habituation the same flies were shifted to and exposed to 10% yeast at 34°C, where the essential function of dynamin in transmitter release would be compromised in Shi^ts1^-expressing cells, then habituation override was significantly impaired as compared to controls (Fig 5A). Thus, yeast-induced dishabituation of control flies not expressing *Shi^ts1^* at 34°C was substantially more efficient (p<0.0001, F-stat = 35.83, Dunn’s multiple comparisons post-hoc test,**p= 0.0015), than of *TH-Gal4, UAS-Shi^ts1^* flies at the same temperature (Fig 5A). Similarly, dishabituation of control flies not expressing *Shi^ts1^* was also substantially more efficient at 34°C (p< 0.0001, F-stat = 46.90, Dunn’s multiple comparisons post-hoc test, ***p = 0.0001), than of *TH-C’-Gal4, UAS-Shi^ts1^* flies, expressing temperature-sensitive mutant dynamin in *THC’-Gal4*neurons. These data argue that presynaptic activity in *TH-C’-Gal4* and *TH-Gal4* positive cells is required for yeast-induced override of PER habituation (Fig 5A).

An important caveat to the above conclusion is that dopaminergic neuron activation may be associated with motivation or reward prediction and therefore be fundamentally required for high levels of PER. In such a scenario, inactivation of TH-expressing cells would be expected to generally reduce levels of PER. To address this issue and test whether the apparently reduced override in Fig 5A is an artifact of blocking dopaminergic neurons during yeast exposure, we measured the innate response to sucrose in naïve flies with and without blockage of synaptic transmission in dopaminergic neurons (Fig 5B). There was no significant difference in the innate response to 2% sucrose, confirming a more restricted role for these TH positive cells in the override of habituation.

### Sensory neurons projections in SEZ overlap spatially with projections of TH-neurons

In the mushroom body, dopaminergic neurons potentially compute novelty by combining excitatory inputs from olfactory sensory channels with inhibitory inputs driven by familiarity encoding interneurons (Zhao et al., 2021) (and see Discussion). If TH-positive cells were to play a similar role in the SEZ, then they would be predicted to receive direct or indirect inputs from taste sensory neurons as well as input from familiarity-representing inhibitory neurons. Previous work has shown that processes from some TH-positive neurons are present in the SEZ which also contains presynaptic endings of taste sensory neurons, processes of local interneurons and dendrites of motor neurons involved in proboscis extension (Figure 6A; (Liu et al., 2012)). Therefore, we considered the possibility that taste sensory neurons make contacts with *TH-*positive processes.

**Figure 6.**
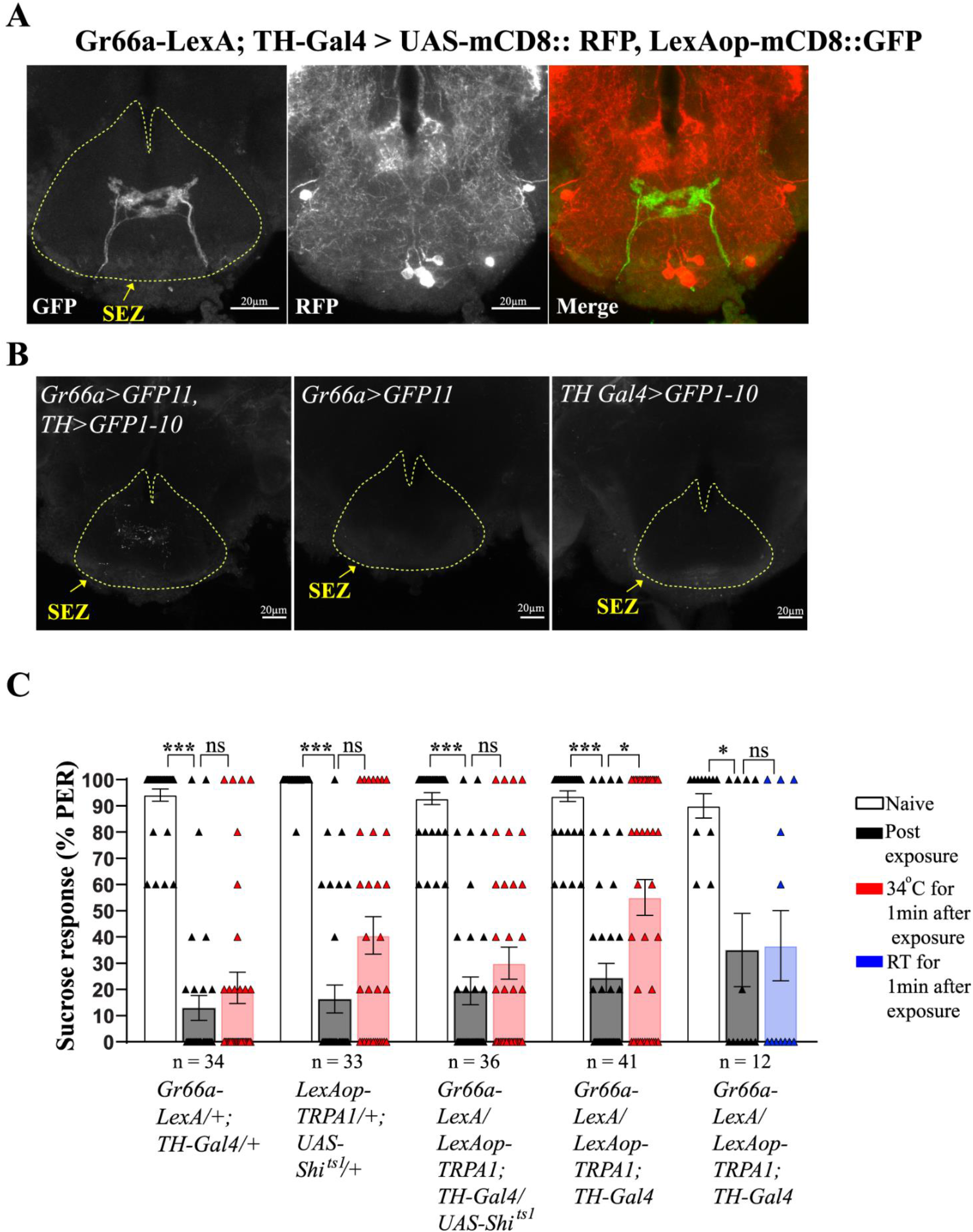
TH Gal4 neurons are functionally connected to GRNs. **(A)** Expression pattern of Gr66a with respect to TH expressing neurons. **(B)** GRASP between Gr66a and TH expressing neurons show GFP signal in animals expressing both the components of split GFP. No signal is observed in controls lacking either of split GFP component. **(C)** Controls do not show any significant increase in response at 34°C (*Gr66aLexA/+; TH-Gal4/+*, p<0.0001, F-statistic = 55.06, Dunn’s multiple comparisons post-hoc test, ***p<0.0001, p= 0.9079, *LexAop-TRPA1/+; UAS-Shi^ts1^/+*, p<0.0001, F-stat = 53.41, Dunn’s multiple comparisons post-hoc test, ***p<0.0001, p = 0.0580,). Activation of Gr66a while inhibiting TH expressing neurons simultaneously fails to reinstate habituated response (*Gr66a LexA/ LexAop-TRPA1; TH-Gal4/UAS-Shi^ts1^*, p<0.0001, F-stat = 52.57, Dunn’s multiple comparisons post-hoc test, ***p<0.0001, p= 0.9494), whereas activating Gr66a solely overrides habituation (*Gr66a LexA/ LexAop-TRPA1; TH-Gal4/ +*, p<0.0001, F-stat = 44.02, Dunn’s multiple comparisons post-hoc test, ***p<0.0001, *p = 0.0205). Control at permissive temperature does not show any difference in habituated response (*Gr66a LexA/ LexAop-TRPA1; TH-Gal4/+*, ***p= 0.0004, F-stat = 15.50, *p= 0.0322, p > 0.99, Dunn’s multiple comparisons post-hoc test). Bars represent mean ± SEM, triangles represent individual data points, ns represents not statistically significantly, p>0.05, SEZ represents subesophageal zone.

To test whether taste-sensory neurons carrying dishabituating signals form connections, direct or indirect, with TH-neuron processes in the SEZ, we used the GFP-reconstitution across synaptic partners (GRASP) technique, which requires the use of dual binary transcription systems (based on LexA and Gal4 transcription factors) to separately express two complementing fragments of GFP, one in sensory neurons and the other in TH neurons. Limited by the immediate availability of transgenes, we were technically restricted to analyzing TH connections with neurons in which gene expression could be controlled by LexA. This was possible for bitter-taste responsive *Gr66a* neurons but not for the yeast-responsive *Ir25a* class.

We first examined whether *Gr66a* axonal projections and *TH-Gal4* marked processes were present in close proximity within the SEZ, by examining the relative localization of GFP driven in *Gr66a*-expressing neurons with RFP in *TH*-positive cells. Processes of sensory neurons expressing *Gr66a-LexA*-driven GFP and dopaminergic neurons expressing *TH-Gal4* driven RFP showed close proximity (Fig6A). Further, GRASP experiments showed that two halves of split GFP, one expressed in *Gr66a* neurons and the other in *TH*-positive cells could combine to reconstitute GFP fluorescence within the SEZ (Fig 6B). Robust fluorescence reconstitution was seen in 8/13 experimental animals compared to 0/15 total animals expressing only one split-GFP component. Thus, GRASP experiments confirm that the processes of *Gr66a* axons and tyrosine-hydroxylase expressing cells come in close proximity of each other. However, due to the limitation of the GRASP technique used it cannot be established certainly if there is a direct synapse between the two cell types.

These data predict that habituation override induced by thermogenetic activation of Gr66a cells should require activity in *TH-Gal4* positive cells. We tested this prediction by creating and analysing PER habituation and dishabituation in *Gr66a-LexA/ LexAop-TRPA1; TH-Gal4/ UAS-Shi^ts1^* and control *Gr66a-LexA/LexAop-TRPA1; TH-Gal4/+* lines at temperatures permissive and restrictive for Shi^ts1^ dynamin. As shown (Fig 6C), both lines showed robust PER and PER habituation. One minute-exposure to 34°C in control flies permissive for synaptic transmission from *TH*-positive neurons resulted in significant (p<0.0001, F-stat = 44.02, Dunn’s multiple comparisons post-hoc test, *p = 0.02) override of habituation. However, similar 34°C thermogenetic stimulation of *Gr66a* neurons in experimental flies where synaptic transmission from TH-positive neurons is blocked did not result in override of PER habituation. Together the anatomical and behavioral data indicate that *Gr66a*-expressing bitter taste sensory neurons are functionally connected to *TH-*expression modulatory neurons whose activity is required for PER habituation override.

Control experiments demonstrating that blocking synaptic output from *TH*-expressing cells has no effect on basal PER response in naïve animals further confirm that transmitter release from *TH*-neurons is required for override of neuronal mechanisms of habituation, not for the innate response to sucrose (Fig 5B).

## DISCUSSION

Animal behavior is profoundly flexible. Thus, at any time, multiple potential behavioral programmes remain dormant, while a subset relevant to specific contexts are active. A growing body of evidence suggests that inhibitory inputs play a major role in preserving perceptions, behaviors and memories in dormant form until required (Barron et al., 2017). Thus, a given context, by recruiting disinhibitory circuits may override inhibition to release latent perceptions, motor programmes and memories appropriate to that context. While the overarching principles are increasingly appreciated, there is still limited understanding of how these are implemented in cells and circuits. While there are many potential reasons for this, the difficulty in studying mechanisms of override has been substantially caused by the paucity of systems in which both mechanisms of habituation or cognitive silencing can be addressed as well as where robust override can be experimentally achieved.

The results we present outline essential elements within a *Drosophila* circuit that overrides habituation of the sucrose-evoked proboscis extension reflex. In doing so, they connect sensory neurons mediating override to neuromodulatory neurons projecting to the SEZ, which may override habituation by silencing inhibitory neurons that drive habituation

Previous work in *Drosophila* concluded that PER habituation arises from increased sucrose-evoked inhibition onto neurons that drive proboscis extension (Paranjpe et al., 2012). Two findings, which closely mirror observations on olfactory habituation, provided key support for this conclusion (Das et al., 2011; Paranjpe et al., 2012). First, the *rutabaga-encoded* adenyl cyclase is required specifically in inhibitory neurons for PER habituation, an observation that we have independently confirmed in this study (Figure 1). Second, experimental silencing of inhibitory neurons causes override of habituation. Together these observations indicate first, that increased GABAergic activity is required for the expression of PER habituation and second, that disinhibition could serve as strategy for habituation override. How might such disinhibition be biologically achieved? Our current experiments show that novel sensory experience induces override of PER habituation through a pathway that requires activity in the *TH-C’* class of dopaminergic neurons. As thermogenetic activation of *TH-C’* cells is also sufficient to override PER habituation, the data suggest a framework in which sensory stimuli activate *TH-C’* neurons, which directly or indirectly, inhibit GABAergic neurons responsible for habituation (Figure 7).

**Figure 7:**
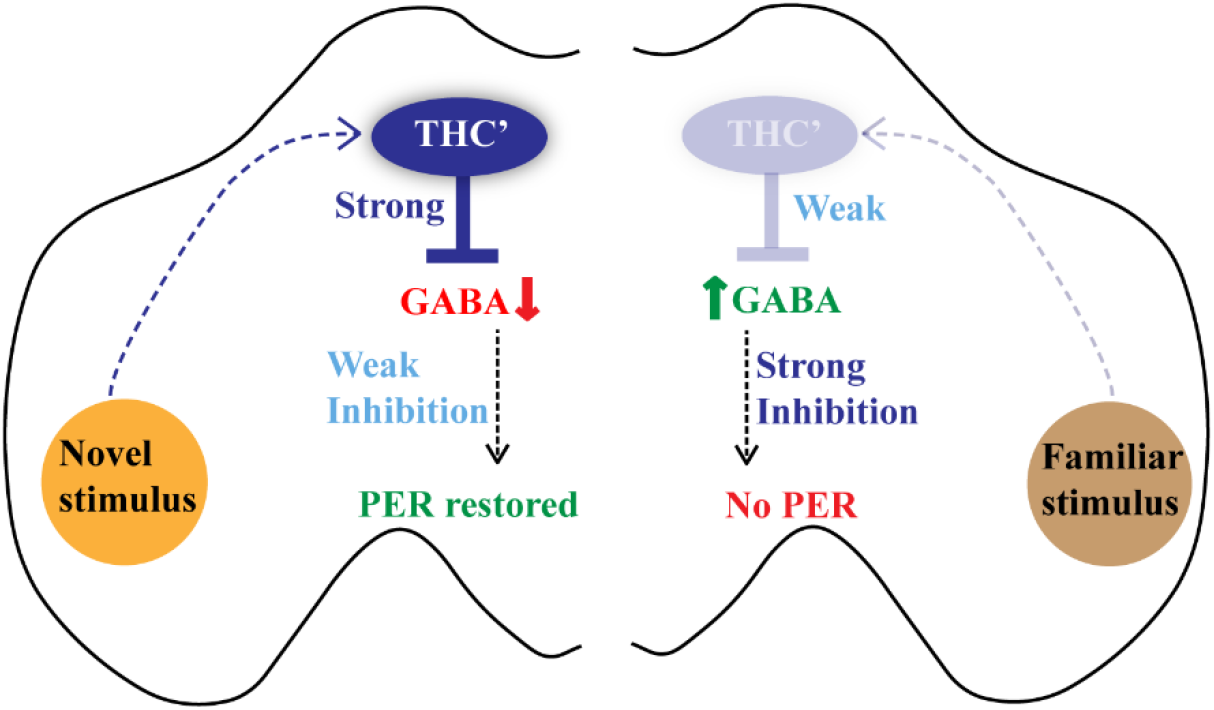
Model for habituation override in the *Drosophila* gustatory system. As shown in Paranjpe et al, 2012, potentiation of GABAergic neurons (“strong inhibition”) underlies PER habituation to a familiar stimulus (Right side). A novel stimulus (left side) activates TH-C’ subset of (likely) dopaminergic neurons which in turn inhibits GABAergic neurons that are potentiated during habituation. The resulting disinhibition restores strong PER to familiar stimulus (habituation override). Familiar stimuli cause only weak activation of THC’ neurons (potentially through increased inhibition onto THC’) and do not alter the habituated PER response.

An important question to address is why novel but not familiar taste stimuli are effective for override? In physiological terms, how could activation of a novel subset of (*Ir25a or Gr66a*) sensory-neurons result in strong excitation of *TH-C’* cells, while similar levels of activation of sucrose-response sensory-neurons causes weaker excitation of the same *TH-C’* cells? We suggest that this occurs because familiar stimuli also result in decreased activation (or likely increased inhibition) of *TH-C’* neurons. A schematic circuit level model is shown in Figure 7. More detailed information on the anatomical and physiological connectivity across key elements of the dishabituation circuit will be required to test and elaborate on this broad model. The subtypes and connectivities of relevant inhibitory neurons in the SEZ remain unknown and crucially important to establish. It is particularly important to know whether different classes of SNs activate different subgroups of iLNs in the SEZ. In addition, the exact subset of *TH-C’* cells involved as well as the mechanism by which they function and modulate GABAergic iLNs need to be identified. While the relevant *TH-C’* cells are marked by three independent dopaminergic reporter lines, *TH-Gal4*, *TH-C’* and *TH-C-Gal80*, and therefore probably dopaminergic, it remains unclear whether dopamine release is required for habituation override. Our attempts to address the last issue via knockdown of tyrosine hydroxylase through RNAi, or by various genetic manipulations of dopamine receptor expression did not yield definitive results (Materials and Methods), often because these manipulations affected baseline levels of PER and PER habituation.

Despite the above lacunae, given that the SEZ, which contains gustatory neurons axons as well as dendrites of motor neurons that drive PER, is likely to be numerically simple, the most parsimonious model for override would posit that taste sensory neurons trigger direct excitation of dopaminergic processes, which in turn acts within the SEZ to directly inhibit GABAergic cells (Pimentel et al., 2016) whose potentiation drives PER habituation.

The model above is consistent with observations on habituation override in mouse and Aplysia brains (Bristol and Carew, 2005; Smith et al., 2009; Kato et al., 2015; Ogg et al., 2018). In mouse, long term auditory habituation to a passively experienced tone is accompanied by increased activity in tone-responsive SOM+ neurons that inhibit similarly tuned pyramidal cells in the auditory cortex. However, if habituated mice are coaxed to attend to the same tone (by a reward for successful engagement), then the behaving mice show overriding inhibition of SOM+ neurons and increased activity of downstream L2/3 pyramidal neurons (Kato et al., 2015). It appears likely that this disinhibition is accomplished by modulatory inputs onto upstream VIP+ neurons. A more recent analysis showed that cholinergic inputs into the mouse olfactory bulb could cause override of a fast form of olfactory habituation. Thus, electrical or optogenetically induced acetylcholine release into the bulb caused mice to override habituation and investigate a previously ignored odor (Ogg et al., 2018). While sensitization is not formally excluded here, these studies are consistent with an emerging theme wherein neuromodulators released in response to novel or meaningful stimuli (Vankov et al., 1995; Giovannini et al., 2001; Ranganath and Rainer, 2003; Hattori et al., 2017; Kafkas and Montaldi, 2018; Morrens et al., 2020) result in disinhibition which can either enhance learning or override habituation.

The experimental results described here provides multiple lines of circumstantial evidence in support of a novelty-induced dopaminergic pathway for disinhibition of sensory perception. It outlines a habituation override circuit all the way from sensory neurons that detect stimulus, to motor neurons that mediate behavioral response. In context of the increasingly widely appreciated role for disinhibition in the control of perception, cognition and behavior (Letzkus et al., 2015; Sridharan and Knudsen, 2015; Barron et al., 2017; Wang and Yang, 2018) we suggest that this work provides a valuable intellectual and biological foundation for future studies to comprehensively identify neurons and mechanisms involved in a central pathway for behaviorally important disinhibition.

## ACKNOWLEDGMENTS

We thank Ali Asgar Bohra, Camilla Roselli, Tamara Boto and James Cooke for comments on the manuscript, Frederic Marion-Poll and members of Ramaswami and VijayRaghavan labs for advice, support and useful discussions. We thank Carlos Ribeiro for valuable discussions and reagents, Pushkar Paranjpe for help setting up the PER assay, as well as the Bloomington stock centre and our many colleagues (mentioned in the text) for various Drosophila stocks used. The work was supported by a Wellcome Trust Investigator Award and a Science Foundation Ireland Investigator Programme Grant to MR, by core support from the National Centre for Biological Sciences (Tata Institute of Fundamental Research) to KVR and by a CSIR postgraduate fellowship and a Biocon-Trinity Scholarship to ST.

## Notes

### Competing Interest Statement

The authors have declared no competing interest.

### Summary of Updates

Introduction updated, Figure 7 revised, Discussion updated to explain the modified model in figure 7.

